# Deep-Learning Based, Automated Segmentation of Macular Edema in Optical Coherence Tomography

**DOI:** 10.1101/135640

**Authors:** Cecilia S. Lee, Ariel J. Tyring, Nicolaas P. Deruyter, Yue Wu, Ariel Rokem, Aaron Y. Lee

## Abstract

Evaluation of clinical images is essential for diagnosis in many specialties and the development of computer vision algorithms to analyze biomedical images will be important. In ophthalmology, optical coherence tomography (OCT) is critical for managing retinal conditions. We developed a convolutional neural network (CNN) that detects intraretinal fluid (IRF) on OCT in a manner indistinguishable from clinicians. Using 1,289 OCT images, the CNN segmented images with a 0.911 cross-validated Dice coefficient, compared with segmentations by experts. Additionally, the agreement between experts and between experts and CNN were similar. Our results reveal that CNN can be trained to perform automated segmentations.

## 1. Introduction

Over the past decade there has been increased interest in deep learning, a promising class of machine learning models that utilizes multiple neural network layers to rapidly extract data in a nonlinear fashion [*1*,*2*]. Trained with large data sets, deep learning has been used successfully for pattern recognition, signal processing and statistical analysis [*2*].Additionally, deep convolutional neural networks (CNN) have facilitated breakthroughs in image processing and segmentation [*1*]. These results have important clinical implications, particularly with regards to interpretation of results from medical imaging procedures. A major advance in computer vision was the development of SegNet, a CNN algorithm that learns segmentations from a training set of segmented images and applies pixel-wise segmentations to novel images[*3*]. Deep learning approaches based on SegNet have been used successfully for automated segmentation in medical imaging problems such as identifying abnormalities in the brain and segmentation of histopathological cells[*4*,*5*]. Although increasing, the application of deep learning has been limited in ophthalmology to detecting retinal abnormalities from fundus photographs [*6*,*7*], glaucoma from perimetry [*8*], segmentation of foveal microvasculature[*9*], and grading of cataracts [*10*].Macular edema, characterized by loss of the blood retinal barrier in retinal microvasculature, leads to accumulation of intraretinal fluid (IRF) and decreased vision [*11*]. With the advent of optical coherence tomography (OCT), the measurement of macular edema inferred by central retinal thickness (CRT) has become one of the most important clinical endpoints in retinal diseases [*12*,*13*]. However, the extent and severity of IRF are not fully captured by CRT [*14*]. Thus, the segmentation of IRF on OCT images would enable more precise quantification of macular edema.Nevertheless, manual segmentations, which require retinal expert identification, are extremetime consuming with variable interpretation, repeatability, and interobserver agreement [*15*,*16*]. Prior studies involving automated OCT segmentations were limited by additional manual input requirements and small validation cohorts [*17*,*18*]. In addition, semiautomated methods included artifact with segmentations [*19*], inaccurately detected microcysts [*20*], and could not distinguish cystic from non-cystic fluid accumulations [*21*]. Recently, we demonstrated that deep learning is effective in classifying OCT images of patients with age-related macular degeneration [*22*]. Using the same OCT EMR database, we sought to apply the CNN for automated IRF segmentations and to validate our deep learning algorithm.

## 2. Materials and Methods

This study was approved by the Institutional Review Board of the University of Washington (UW) and was in adherence with the tenets of the Declaration of Helsinki and the Health Insurance Portability and Accountability Act.Macular OCT scans from the period 2006-2016 were extracted using an automated extraction tool from the Heidelberg Spectralis imaging database at the University of Washington Ophthalmology Department. All scans were obtained using a 61-line raster macula scan, and every image of each macular OCT was extracted. Clinical variables were extracted in an automated fashion from the Epic electronic medical records database in tandem with macular OCT volumes from patients that were identified as relevant to the study, because they had received therapy for diabetic macular edema, retinal vein occlusions, and/or macular degeneration. The central slice from these OCT volumes was selected for manual segmentation.

A custom browser-based application was implemented (using HTML5) and a webpage was created to passively record the paths drawn by clinicians when segmenting the images. IRF was defined as ‘an intraretinal hyporeflective space surrounded by reflective septae” as used in other studies[*17*,*23*]. Each segmentation result was then reviewed by an independent retina-trained clinician and inaccurate segmentations were rejected. These images were then divided: 70% were designated as a *training set* used for optimization of the CNN. The remaining 30% were designated as a *validation set*. These were used for crossvalidation during the optimization procedure. An additional final *test set* of 30 images were selected, ensuring that none of the patients from the training and validation sets were present in the test set. The images in the *test set* were segmented by four independent clinicians. Since retinal OCTs images have variable widths, a vertical section of 432 pixels tall and 32 pixels wide was chosen *a priori* with the plan to slide the window across the images and collect the predicted probabilities from the CNN. The outputted probabilities for each pixel were then averaged to create a probability distribution map.

In the training set, data augmentation was performed by varying the position of the window on the OCT image. Care was taken in the validation set to ensure that no overlap was present in any of the validation images. In addition, the training and validation sets were balanced such that 50% of the images contained at least one pixel marked as intraretinal fluid while the other half were from images without any intraretinal fluid.

A smoothed Dice coefficient was used as the loss function and as the assessment of the final held-out test set with the smoothness parameter set to 1.0.

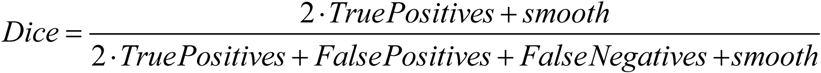

The CNN used was a modified version of the U-net autoencoder architecture[*24*], with a total of 18 convolutional layers (Figure 1). The final activation function was a sigmoid function that generates a prediction of a binary segmentation map.

**Fig. 1.**
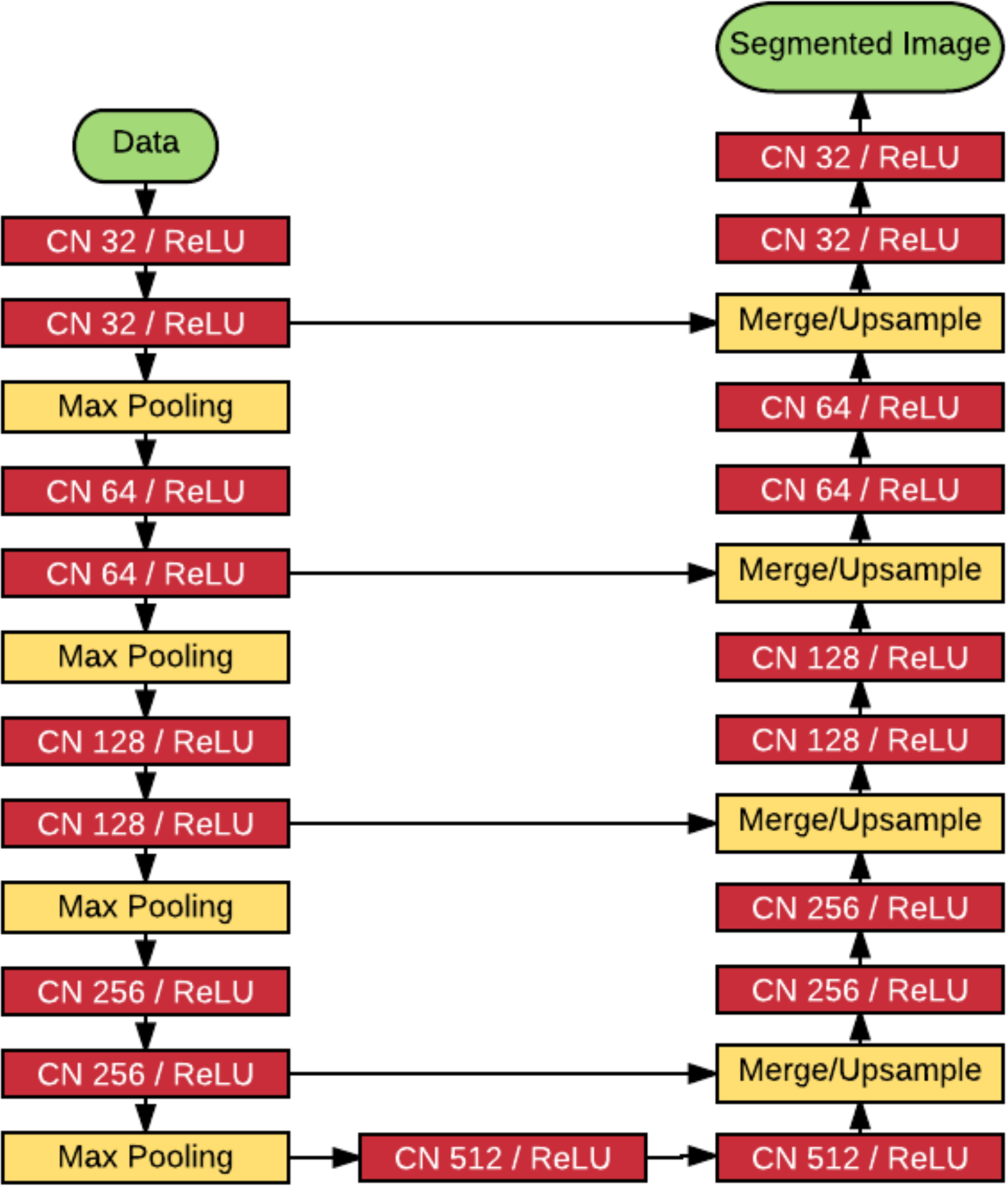
Schematic of deep learning module using a total of 18 convolutional layers. CNN, convolutional neural network. ReLU, rectified linear unit.

The model was trained using the Adam optimizer with a learning rate set at 1e-7. All training occurred using Keras (https://keras.io) and Tensorflow (arXiv:1603.04467) using NVIDIA Tesla P100 Graphics Processing Units (GPU) with NVIDA cuda (v8.0) and cu-dnn (v5.5.1) libraries (http://www.nvidia.com). Statistics were performed using R (http://www.r-project.org). The model, along with the trained final weights and the code to run the model,have been published as an open source project: (https://github.com/uw-biomedical-ml/irfsegmenter).

In assessing the final held-out test set, the CNN performed inference using a sliding window generating an array of probabilities for each pixel. Then this output was compared using the above Dice coefficient to baseline segmentations performed by clinicians.

In addition, the CNN was compared against four clinicians. For this comparison, an individual clinician was chosen as the groundtruth, and the Dice coefficient of the other three clinicians as well as the CNN were computed from this groundtruth. This was repeated for all four clinicians so that Dice coefficients with each clinician as baseline were computed. The coupled Dice coefficients showed no statistical difference between the clinicians and the CNN.

## 3. Results

Manual segmentations of 1,289 OCT macular images were used for training and cross validation. A total of 1919680 sections of 934 manually segmented OCT images were designated as a training set: these were presented to the CNN, and the segmentation errors from these images were used to adjust the weights in different layers of the networks. Another 1,088 sections of 355 images were used for cross-validation: they were presented to the CNN during training and segmentation errors were recorded, but were not used to adjust the weights in the network layers The model was trained with 200,000 iterations and periodically assessed against the validation set (cross-validation). The learning curve showing the Dice coefficients of the training iterations and the validation set are shown in Figure 2. The model achieved a maximal cross-validation Dice coefficient of 0.911.

**Fig. 2.**
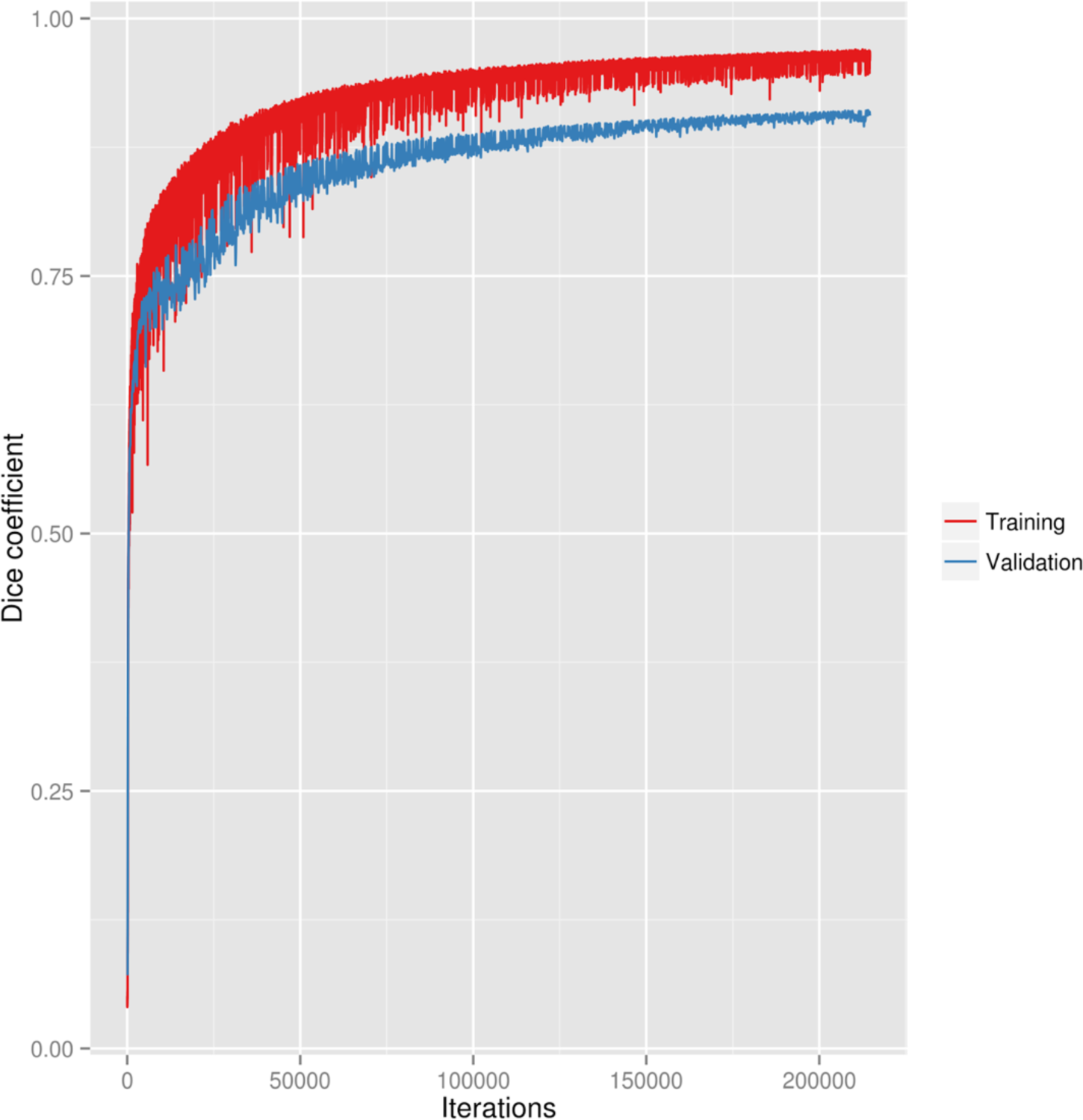
Learning curve of the neural network training and validation.

The weights that produced the highest cross-validated coefficient were then used for comparison against manual segmentations in a held out test set. Example deep learning segmentations of intraretinal fluid from the held out test set are shown in Figure 3.

**Fig. 3.**
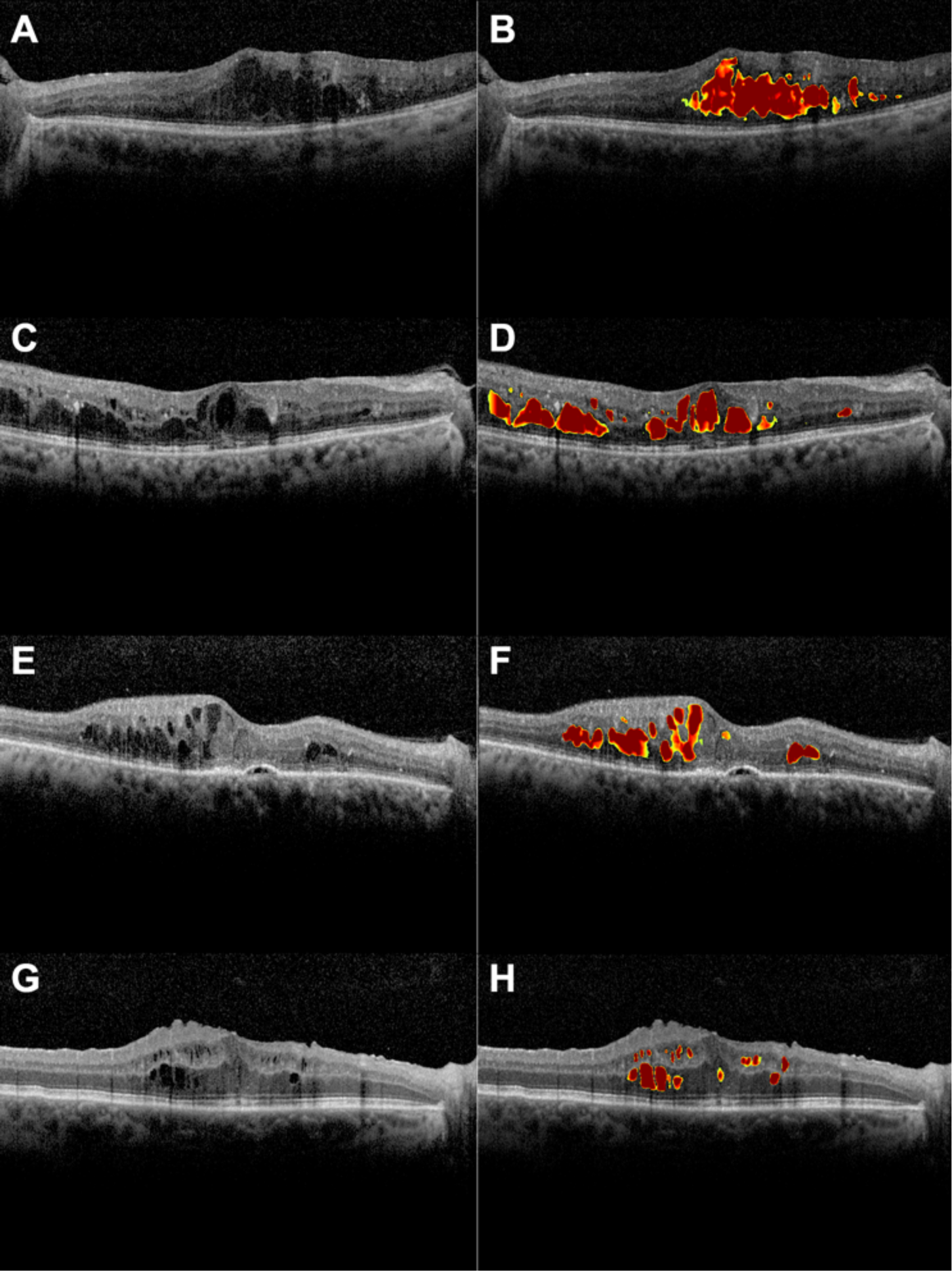
Example segmentations of intraretinal fluid from the held out test set by deep learning. A, an example optical coherence tomography (OCT) image with intraretinal fluid. B, automated segmentation by deep learning. C, an example optical coherence tomography (OCT) image with intraretinal fluid. D, automated segmentation by deep learning. E, an example OCT image with intraretinal and subretinal fluid. F, deep learning correctly segments intraretinal fluid cysts but not subretinal fluid. G, an example optical coherence tomography (OCT) image with intraretinal fluid and an epiretinal membrane with macular pucker. H, deep learning correctly segments intraretinal fluid cysts but not sub-internal limiting membrane spaces.

To assess reliability of the deep learning algorithm we compared the Dice coefficientsfordeeplearning automated segmentations against manual segmentations by four independent clinicians. The pairwise comparison of Dice coefficients are shown in Table 1.

**Table 1.** Mean dice coefficients with standard deviations.

|  | Clinician 2 | Clinician 3 | Clinician 4 | Deep Learning |
| --- | --- | --- | --- | --- |
| Clinician 1 (N) | 0.744 (0.185) | 0.771 (0.155) | 0.809 (0.164) | 0.754 (0.135) |
| Clinician 2 (C) |  | 0.710 (0.190) | 0.747 (0.189) | 0.703 (0.182) |
| Clinician 3 (A) |  |  | 0.722 (0.209) | 0.725 (0.117) |
| Clinician 4 (Ar) |  |  |  | 0.730 (0.176) |

**The clinician in the first column is the baseline from whom Dice coefficients are computed for each row.**

The mean Dice coefficient for human interrater reliability and deep learning were 0.750 and 0.729, respectively, reaching a generally accepted value of excellent agreement (Figure 4) [*25*]. No statistically significant difference was found between clinicians and deep learning (p = 0.247).

**Fig. 4.**
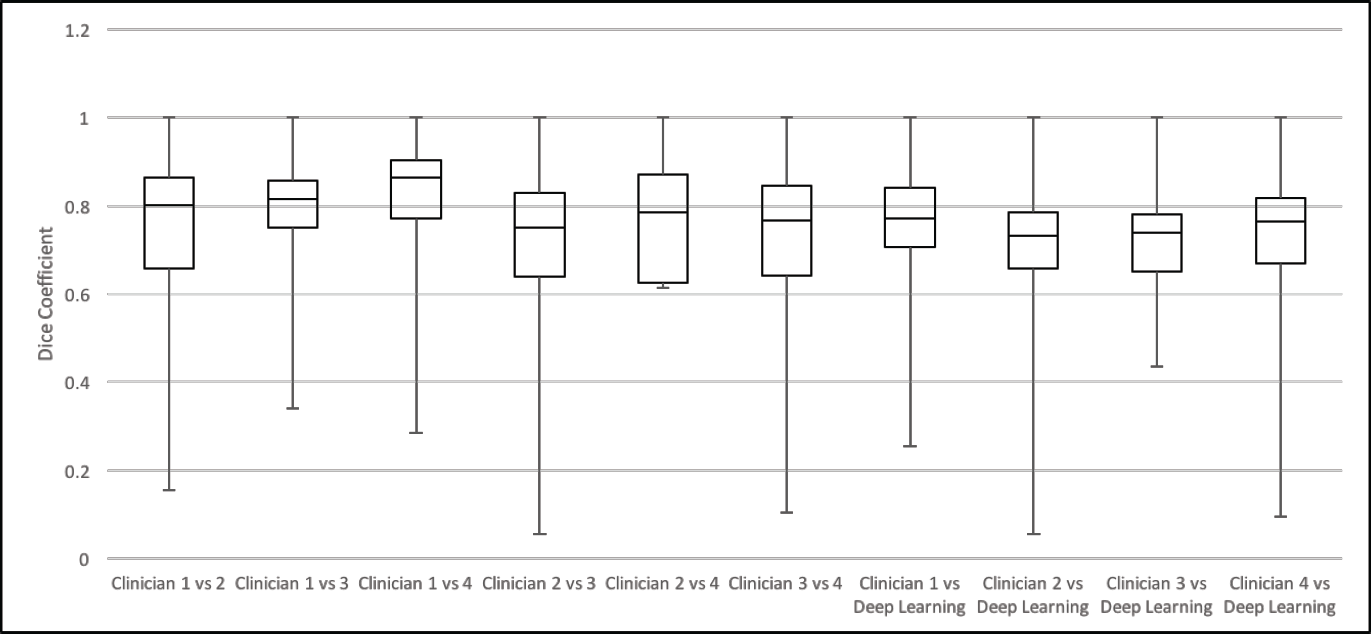
Pairwise comparison of Dice coefficients from the held out test set. The mean Dice coefficient for human interrater reliability and deep learning were 0.750 and 0.729, respectively.

## 4. Discussion

The concept of automated segmentation using deep learning has become increasingly popular in medical imaging; however, comparatively little has been applied in ophthalmology and the validation of proposed models were limited to small sample size. Zheng et al. evaluated semiautomated segmentation of intra- and subretinal fluid in OCT images of 37 patients [*17*].Initial automated segmentation was performed followed by coarse and fine segmentation of fluid regions. Then an expert manually selected each region of potential IRF and SRF. This semi-automated segmentation had a Dice coefficient of 0.721 to 0.785 compared to manual segmentation, which is similar to our study. The advantage of our algorithm includes being a fully automated system with no human input. Esmaeili et al. used recursive Gaussian filter and 3D fast curvelet transform to determine curvelet coefficients, which were denoised with a 3D sparse dictionary. The average Dice coefficients for the segmentation of intraretinal cysts in whole 3D volume and within central 3 mm diameter were 0.65 and 0.77, respectively [*26*]. However, the validation of their study included only four 3D-OCT images, a much smaller sample than our study. Chiu et al. reported a dice coefficient of 0.53 using an automated detection of IRF in OCT images of 6 patients with DME. Both Dice coefficient and intermanual segmenter correlation (0.58) were much lower than our study [*21*].

Our study shows that CNN can be trained to identify clinical features that are diagnostic, similar to how clinicians interpret an OCT image. The deep learning algorithm successfully identified areas of IRF by generating pixelwise probabilities which were then averaged to create a segmentation map. Our model was trained with more than 200,000 iterations without overfitting. Additional advantages of this study include the use of a publicly available model and training strategy for precise localization and segmentation of images.Furthermore, the deep learning model is fast and requires no additional human input, such as with semi-automated segmentation programs. To our knowledge, the use of a validated CNN for accurate segmentation of IRF has never been shown.

Our training datasets originated from images from a single academic center, thus our results may not be generalizable. However, our training dataset was randomly selected from all consecutive retinal OCT images obtained at our institution during a 10-year period, therefore a wide variety of pathologies likely have been included, increasing its overall applicability in other datasets. In summary, deep learning shows promising results in accurate segmentation and automated quantification of intraretinal fluid volume on OCT imaging and our open-source algorithm may be useful for future clinical and research applications

## 5. Funding and Acknowledgements

National Eye Institute, Bethesda, Maryland (grant no.: K23EY02492 [C.S.L.]); Latham Vision Science Innovation Grant, Seattle WA (C.S.L.); Research to Prevent Blindness, Inc., New York, New York (C.S.L., A.Y.L.); The Gordon & Betty Moore Foundation and the Alfred P. Sloan Foundation University of Washington eScience Institute Data Science Environment (A.R.).

